# Mitochondrial DNA Variation in the D-LOOP and ND Loci identified in the Kenyan Population: Potential Implications for precision Oncology

**DOI:** 10.64898/2026.03.12.711313

**Authors:** Eva A Nambati, Belinda Azzam, Eunice Githua, Ngure Kirosh, Lydia S. Eyase, Solomon Langat, Sharon Ariga, Winfrida Cheriro, Fredrick Eyase, Wallace Bulimo

## Abstract

**Background:** Precision oncology is dominated by studies focused on nuclear genomic alterations, leaving mitochondrial DNA (mtDNA) variation excluded from routine clinical genomic testing. However, mitochondria regulate oxidative phosphorylation (OXPHOS), reactive oxygen species (ROS) production, apoptosis, and metabolic reprogramming pathways that are central to chemotherapy response.

**Methods:** 468 Complete mitochondrial genomes from Kenyan individuals representing diverse ethnic groups were analyzed. Seven variants associated with effect on cancer treatment were identified. These include; m.310T>C(D-loop), m.10398A>G (MT-ND3), m.13708G>A (MT-ND5), m.16189T>C, m.13928G>C, m9055G>A and m.16519T>C (D-loop). Allele frequencies and distribution were assessed.

**Results:** The coding-region variants (m.10398A>G and m.13708G>A) occur in Complex I subunits and are associated with altered oxidative phosphorylation efficiency and ROS production. The control-region variants (m.16189T>C and m.16519T>C) influence mtDNA replication and copy number. These variants have been implicated in differential response to chemotherapeutic agents including platinum-based therapies and anthracyclines. m.13928G>C sits in the MT-CYB gene and could possibly affect mitochondrial respiratory function; this variant could influence how tumors respond to therapies that rely on apoptosis or ROS generation.m.9055G>A is a MT-ATP6 variant classified as benign in mitochondrial disease but may represent a marker of haplogroup background rather than a direct cancer driver. While m.310T>C itself does not encode a protein, its location in the regulatory D-loop influences mitochondrial function, which can affect how tumor cells respond to chemotherapies that rely on mitochondrial-mediated apoptosis or oxidative stress.

**Conclusion:** Pharmacogenomic relevant mitochondrial variants are present in the Kenyan population. With the rise of cancer burden in Kenya there is a need carry out more studies to understand the impact of these variations on cancer treatment. This can inform the integration of mtDNA analysis into precision oncology strategies in African populations.

## Introduction

Precision oncology primarily relates to nuclear genomic alterations, that aid precise diagnosis and guide personalized treatments. Mitochondria play vital roles in cellular bioenergetics, reactive oxygen species (ROS) production and regulation of intrinsic apoptosis. All these processes are directly relevant to cancer development and treatment response, yet its genome is seldom considered in precision oncology (Vyas et al., 2016; Wallace, 2012). Many chemotherapeutic agents induce cell death by promoting oxidative stress or activating mitochondrial apoptotic pathways. Hence inherited variations (germline mutations) or acquired (somatic mutations) in the mitogenome can either enhance cancer cells death or aid resistance (Khair et al., 2024; Wallace, 2012)

### Mitochondrial DNA Variation and role in Cancer

The mitochondrial displacement loop (D-loop) is a non-coding control region that plays a critical role in regulating mitochondrial DNA (mtDNA) replication and transcription. Due to its lack of protective histones and limited DNA repair mechanisms, the D-loop exhibits a higher mutation rate than other regions of the mitochondrial genome(Masuda et al., 2012; Sharma et al., 2005) (Masuda et al., 2012; Sharma et al., 2005). Numerous studies have reported somatic mutations and polymorphisms within the D-loop in a wide range of cancers, including breast, colorectal, hepatocellular, and gastric cancers(Lin et al., 2015) (Lin et al., 2015). D-loop mutations can disrupt mitochondrial homeostasis, leading to increased production of reactive oxygen species (ROS), genomic instability, and metabolic reprogramming that supports tumor growth and progression(Lee & Wei, 2009; Lu et al., 2009; Shimura & Kunugita, 2016)(Lee & Wei, 2009; Lu et al., 2009; Shimura & Kunugita, 2016), because of its high variability and frequent alteration in tumors, the D-loop region has also been investigated as a potential biomarker for cancer detection, prognosis, and monitoring of disease progression.(Czarnecka et al., 2010; Harino et al., 2024a; Vega Avalos et al., 2022)(Czarnecka et al., 2010; Harino et al., 2024a; Vega Avalos et al., 2022)

Genes encoding the NADH dehydrogenase (ND) subunits of mitochondrial complex I play a central role in oxidative phosphorylation and cellular energy production. Variations within ND genes (ND1–ND6 and ND4L) have been widely implicated in cancer development and progression due to their impact on mitochondrial respiratory function. Mutations in these genes can impair electron transport chain activity, resulting in altered ATP production and increased generation of reactive oxygen species, which may promote oxidative stress and tumorigenesis(Brandon et al., 2006; Wallace, 2012). Additionally, dysfunction of complex I caused by ND mutations can induce metabolic shifts toward glycolysis, a hallmark of cancer cell metabolism that facilitates survival under hypoxic conditions and supports rapid cell proliferation(Sollazzo et al., 2022). Several studies have identified recurrent ND gene mutations in cancers such as prostate, colorectal, pancreatic, and lung cancers, suggesting their potential role in tumor progression and therapeutic response(Yuan et al., 2020).

The MT-CYB encodes a key component of mitochondrial complex III in the electron transport chain, which is essential for oxidative phosphorylation and cellular energy production. Variations in MT-CYB can disrupt electron transport efficiency, leading to altered production of reactive oxygen species (ROS) and changes in mitochondrial membrane potential. These alterations may promote tumor development by increasing oxidative stress, genomic instability, and metabolic reprogramming in cancer cells(Wallace, 2012).

The MT-ATP6 encodes a subunit of ATP synthase (complex V), the enzyme responsible for ATP production during oxidative phosphorylation in mitochondria. Variations in MT-ATP6 can affect proton transport across the mitochondrial membrane and alter the efficiency of ATP synthesis. Such disruptions may influence cellular energy metabolism, a process frequently reprogrammed in cancer cells to support rapid proliferation and survival(Jonckheere et al., 2012; Wallace, 2012). Since mitochondria play a central role in regulating cell death, variants in MT-ATP6 may influence tumor cell survival and potentially affect responses to therapies that target mitochondrial metabolism or induce apoptosis. Although many MT-ATP6 variants are classified as benign in mitochondrial disease, their presence in tumors may still contribute to metabolic variability and cancer cell adaptation(Vyas et al., 2016).

## Methods

468 mitochondrial genomes from Kenyan individuals representing diverse ethnolinguistic groups were analyzed. Using a bash script, paired-end FASTQ files were aligned with BWA-MEM and converted from SAM to sorted, indexed BAM files using Samtools. The MutServe variant caller was used, Variants were detected by comparing reads against the revised Cambridge Reference Sequence (rCRS). The mitochondrial reference genome used was NC_012920.1.

Variant annotation was carried out to determine the location, effect, and potential clinical significance.

Seven variants associated with cancer were identified: m.310T>C(D-loop), m.10398A>G (MT-ND3), m.13708G>A (MT-ND5), m.16189T>C, m.13928G>C, m9055G>A and m.16519T>C (D-loop). Allele frequencies and distribution were assessed.

## Results

A total of 2,278 variants were identified in the study, of which 350 (15%; 350/2,278) were mitochondrial variants associated with cancer. Table 1 below.

**Table 1:**
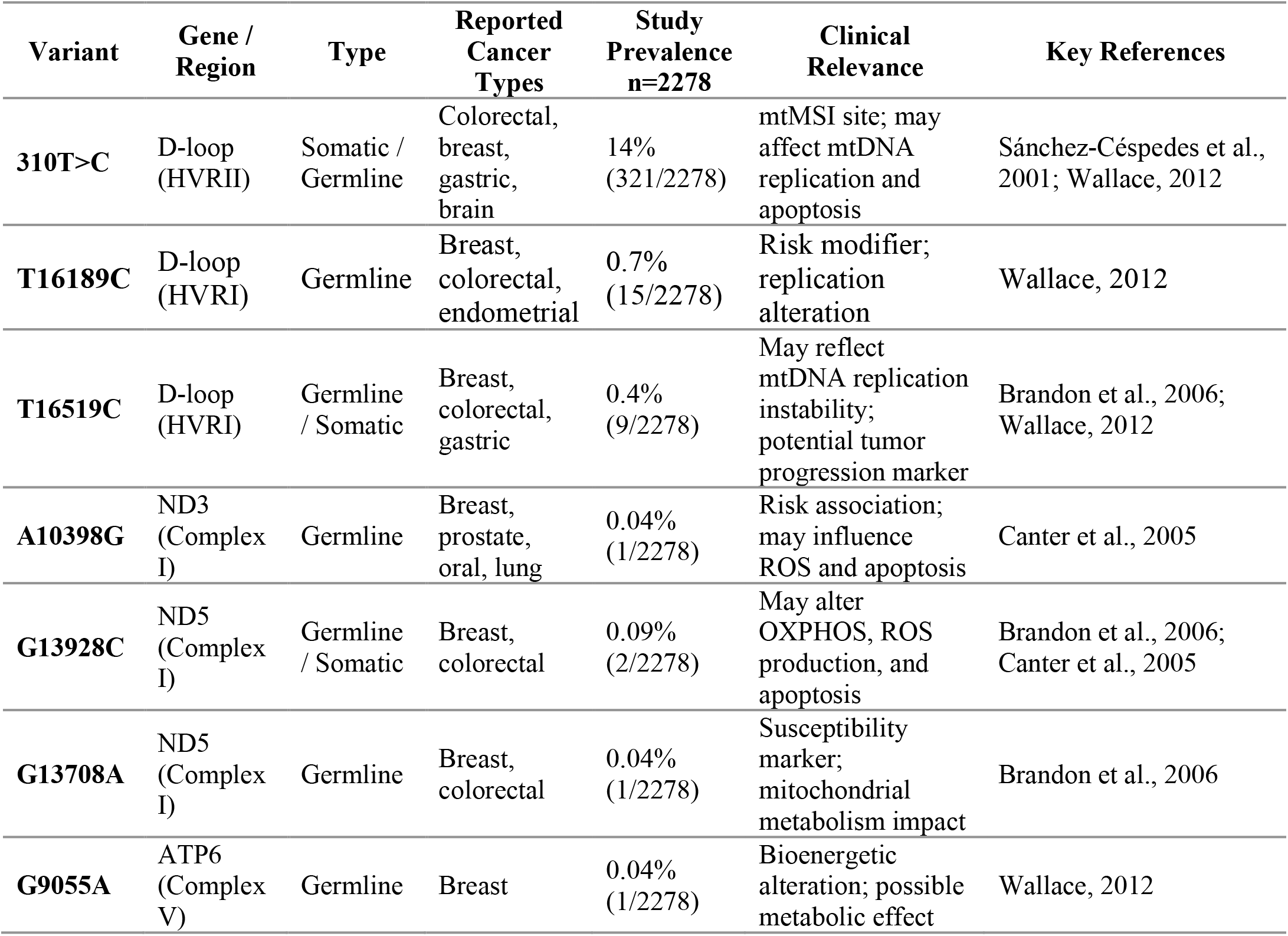
Identified Human mtDNA Variant Cancer Markers and their clinical implications.

| Variant | Gene / Region | Type | Reported Cancer Types | Study Prevalence n=2278 | Clinical Relevance | Key References |
| --- | --- | --- | --- | --- | --- | --- |
| **310T>C** | D-loop (HVRII) | Somatic / Germline | Colorectal, breast, gastric, brain | 14% (321/2278) | mtMSI site; may affect mtDNA replication and apoptosis | Sánchez-Céspedes et al., 2001; Wallace, 2012 |
| **T16189C** | D-loop (HVRI) | Germline | Breast, colorectal, endometrial | 0.7% (15/2278) | Risk modifier; replication alteration | Wallace, 2012 |
| **T16519C** | D-loop (HVRI) | Germline / Somatic | Breast, colorectal, gastric | 0.4% (9/2278) | May reflect mtDNA replication instability; potential tumor progression marker | Brandon et al., 2006; Wallace, 2012 |
| **A10398G** | ND3 (Complex I) | Germline | Breast, prostate, oral, lung | 0.04% (1/2278) | Risk association; may influence ROS and apoptosis | Canter et al., 2005 |
| **G13928C** | ND5 (Complex I) | Germline / Somatic | Breast, colorectal | 0.09% (2/2278) | May alter OXPHOS, ROS production, and apoptosis | Brandon et al., 2006; Canter et al., 2005 |
| **G13708A** | ND5 (Complex I) | Germline | Breast, colorectal | 0.04% (1/2278) | Susceptibility marker; mitochondrial metabolism impact | Brandon et al., 2006 |
| **G9055A** | ATP6 (Complex V) | Germline | Breast | 0.04% (1/2278) | Bioenergetic alteration; possible metabolic effect | Wallace, 2012 |

Seven variants associated with cancer were identified: m.310T>C(D-loop), m.10398A>G (MT-ND3), m.13708G>A (MT-ND5), m.16189T>C, m.13928G>C, m9055G>A and m.16519T>C (D-loop). Allele frequencies and distribution were assessed as shown in Fig 1 below.

**Fig. 1:**
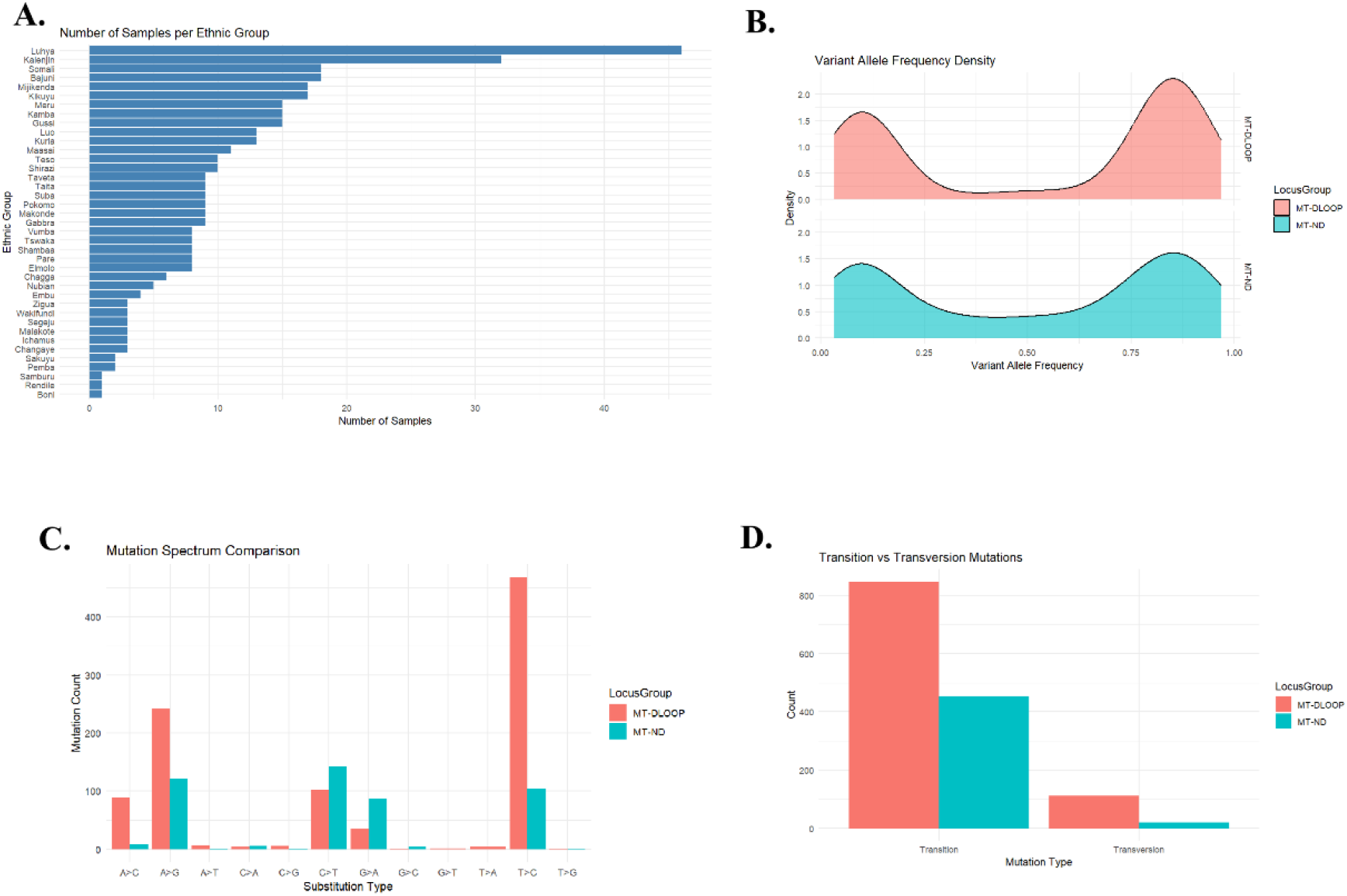
A. Samples analysed - per ethnic group, B. Variant allele frequency density plot, C. Mutation spectrum comparison, D. Transition vs. Transversion (Ti/Tv) Plot

### Mitochondrial DNA Variation in the D-LOOP and ND Loci in the Kenyan Population

Although variants were present in other regions such as the MT-ATP6. The analysis focused on genetic variations within the D-LOOP and ND loci in different individuals from Kenya where frequency was relatively higher. The samples used in this study were derived from ethnically diverse groups from different geographical regions in Kenya. The Luhya and Kalenjin represented the largest sample groups, followed by the Somali, Bajuni, and Mijikenda. Smaller sample sizes were obtained from groups such as the Boni, Rendile, and Samburu. Notably, the variants were identified across all the groups.

The distribution of Variant Allele Frequency (VAF) indicates the presence of both low-frequency and high-frequency variants in both studied loci. Both the MT-DLOOP and MT-ND regions exhibit a bimodal distribution of variant allele frequencies. Peaks are observed at low VAF (approx. 0.1) and high VAF (approx. 0.8–0.9).

Distinct mutational signatures were identified in both the D-LOOP and ND genes. The T > C substitution was the most prevalent mutation type in the MT-DLOOP region, followed by A > G. While the T > C was also frequent in the MT-ND region, its count was significantly lower than in the D-LOOP. Conversely, the C > T substitution showed a relatively higher frequency in the MT-ND locus compared to other types within that same region.

Transition Mutations were significantly more frequent than transversions in both regions. MT-DLOOP exhibited a substantially higher total count of transitions (over 800) compared to the MT-ND region (approx. 450). Transversion Mutations occurred at much lower frequencies, with the MT-DLOOP again showing a higher count of transversions relative to the MT-ND region.

## Discussion

The observed varying Variant Allele Frequency suggests varying levels of heteroplasmy. High-frequency variants (VAF > 0.8) likely represent fixed or near-fixed mutations in maternal lineages across the Kenyan ethnic groups.

This disparity in the frequency of variants is expected given that the D-LOOP is a non-coding control region with fewer evolutionary constraints than the protein-coding ND genes. The high frequency of T > C and A > G transitions in the D-LOOP may reflect a localized hypermutability that could serve as a population-specific genetic marker.

A strong bias toward transition mutations was observed across both loci, which is characteristic of mitochondrial DNA evolution.

Mitochondrial DNA (mtDNA) alterations have been widely investigated for their potential role in cancer development and progression and pharmacogenomic effects. Both somatic mutations and inherited polymorphisms have been reported in tumor tissues, with some variants proposed as diagnostic, prognostic, or susceptibility markers. However, the strength of these associations varies across populations and cancer types(Wallace, 2012).

One of the most frequently studied regions of mtDNA in cancer research is the displacement loop (D-loop), particularly the D310 mononucleotide repeat located in hypervariable region II (HVRII). This polycytosine stretch is considered a mutational hotspot and is commonly altered in colorectal, breast, gastric, liver, and brain tumors (Ka et al., 2024; Zhu et al., 2005). Because D310 instability is often detected in early-stage tumors, it has been proposed as a potential biomarker for early cancer detection. Another D-loop variant, D16184C in hypervariable region I (HVRI), has been reported in gastric and endometrial cancers and is thought to influence mtDNA replication stability (Brandon et al., 2006) (Brandon et al., 2006). Similarly, the T16189C polymorphism has been examined in breast, colorectal, and endometrial cancers. Although typically considered a population polymorphism, it has been suggested to modify mitochondrial replication efficiency and potentially influence cancer susceptibility (Wallace, 2012) (Wallace, 2012).

Variants in genes encoding components of the oxidative phosphorylation (OXPHOS) system, particularly complex I, have also received significant attention. The A10398G polymorphism in the ND3 gene is one of the most extensively studied mtDNA variants in cancer. It has been investigated in breast, prostate, oral, and lung cancers, with some studies reporting increased risk in specific ethnic groups(Canter et al., 2005) (Canter et al., 2005). However, findings have not been consistent across populations. Other complex I variants, such as G13708A in the ND5 gene, have also been reported in breast and colorectal cancers, though their functional significance remains under evaluation (Brandon et al., 2006) (Brandon et al., 2006).

Transfer RNA (tRNA) gene variants represent another category of mtDNA changes studied in oncology. The A12308G polymorphism in the tRNALeu (CUN) gene has been described in breast, colorectal, prostate, and renal cancers. Some haplotype-based studies have suggested that it may act as a cancer susceptibility marker in certain populations (Covarrubias et al., 2008) (Covarrubias et al., 2008). Another variant, G15927A in the tRNAThr gene, has been observed in gastric and colorectal tumors, although its clinical relevance requires further clarification (Brandon et al., 2006) (Brandon et al., 2006). The well-known pathogenic mutation A3243G in tRNALeu (UUR), primarily associated with mitochondrial disorders, has also been detected in various solid tumors, suggesting that severe mitochondrial dysfunction may contribute to tumor biology (Wallace, 2012) (Wallace, 2012).

Alterations in other components of the respiratory chain have also been described. Variants in the cytochrome b (CYB) gene, such as G15059A and T14798C, have been reported in prostate, bladder, and breast cancers (Brandon et al., 2006) (Brandon et al., 2006). Additionally, mutations in ATP synthase genes, including G9055A and the pathogenic T8993G mutation in ATP6, have been identified in tumor tissues and may influence cellular energy metabolism (Wallace, 2012) (Wallace, 2012). Beyond point mutations, large-scale mtDNA deletions, particularly the common 4977 bp deletion, have been detected in breast, gastric, and liver cancers. These deletions are often linked to oxidative stress and mitochondrial damage (Brandon et al., 2006)

Overall, numerous mtDNA variants have been proposed as cancer-associated markers. While some alterations, especially those in the D-loop region, appear frequently in tumors, many reported associations are population-specific or remain controversial. Consequently, although mitochondrial variants show promise as biomarkers, further large-scale and well-controlled studies are necessary to establish their definitive clinical utility.

Mitochondrial genetic variation does not typically alter drug metabolism directly in the way that nuclear-encoded cytochrome P450 enzymes do. Instead, mitochondrial variants function as pharmacogenomic modifiers by influencing intracellular pathways that determine how cancer cells respond to therapeutic stress(Harino et al., 2024) (Harino et al., 2024). Because mitochondria regulate oxidative phosphorylation, reactive oxygen species (ROS) generation, and intrinsic apoptotic signaling, alterations in mtDNA can shift cellular redox balance, modify apoptotic thresholds, and affect metabolic adaptability under chemotherapeutic pressure (Vyas et al., 2016; Wallace, 2012) (Vyas et al., 2016; Wallace, 2012). These mechanisms suggest that mitochondrial background may contribute to variability in cytotoxic drug sensitivity and resistance.

Increasing evidence indicates that mtDNA variation can influence treatment response through modulation of ROS-mediated cytotoxicity and mitochondrial-dependent apoptosis (Brandon et al., 2006; Wallace, 2012) (Brandon et al., 2006; Wallace, 2012). Since many anticancer agents partly exert their effects by inducing oxidative stress or triggering mitochondrial apoptosis, interindividual differences in mitochondrial genome composition may alter therapeutic outcomes, Thus, mtDNA variants may act as indirect determinants of chemotherapy efficacy rather than classical pharmacokinetic regulators(Ju et al., 2024) (Ju et al., 2024).

In African populations, including those in Kenya, high levels of mitochondrial genetic diversity have been documented due to deep ancestral lineages and extensive haplogroup variation(Gomez et al., 2014) (Gomez et al., 2014). Given this diversity, mitochondrial genomic differences may represent an underrecognized contributor to variability in chemotherapy response across individuals. However, pharmacogenomic studies in Africa remain comparatively limited, and mitochondrial variation is rarely incorporated into oncology research frameworks (Yalcouyé et al., 2026) (Yalcouyé et al., 2026). This gap may partially contribute to inconsistencies in treatment outcomes and observed disparities in cancer survival across regions.

Integrating mitochondrial genome analysis into oncology testing strategies in African settings could therefore enhance risk stratification and individualized treatment planning. Expanding precision oncology efforts to include mtDNA sequencing alongside established nuclear pharmacogenomic markers may help clarify population-specific response patterns and reduce inequities in cancer care (Dandara et al., 2019). Further large-scale, well-controlled studies are needed to determine the clinical utility of mitochondrial variants in predicting chemotherapy response within diverse African populations.

Because many cancer therapies, including chemotherapeutics and targeted inhibitors induce oxidative stress or mitochondrial-mediated apoptosis, the mitochondrial variants of a patient may determine their response to treatment. Mutations in the ND genes can lead to altered Reactive Oxygen Species (ROS) production, potentially providing a survival advantage to malignant cells and contributing to therapy resistance.

## Conclusion

Mitochondrial DNA variants with functional relevance to oxidative phosphorylation and apoptosis are prevalent in the Kenyan population. This can potentially affect cancer treatment and prognosis. Incorporating mtDNA analysis into precision oncology therefore has the potential to improve personalized cancer care in African populations.

D310 and D16184 are among the best-characterized mitochondrial microsatellite loci associated with cancer. From our analysis D310 and T16189C were the most prevalent, however further studies are needed to clarify the clinical significance. While mtDNA variants such as A12308G and A10398G have been widely investigated in cancer research, their exact functional roles and clinical utility remain subjects of ongoing debate. The A10398G variant was observed in this study.

As we advance in precision oncology it will be worthwhile to investigate mitochondrial variants in cancer patients and correlate these variants with clinical treatment outcomes in Kenya.

